# Evaluation of visual performance for RCS rats: A tailor-made OMR setup via DeepLabCut and Psychopy with a customized algorithm

**DOI:** 10.1101/2025.05.24.655932

**Authors:** Jing Wang, Thomas Rüland, Sandra Krause, Michael Schöneck, N. Jon Shah, Frank Müller, Björn Kampa, Karl-Josef Langen, Antje Willuweit

## Abstract

In this study, we developed an optimized optomotor response (OMR) setup for rats based on existing commercial systems and introduced a novel method that integrates DeepLabCut, PsychoPy, and custom algorithms to automate OMR performance evaluation in rats. Our primary findings revealed the optimal spatial frequency (SF) and grating velocity for pigmented Royal College of Surgeons (RCS) and wild-type (WT) rats. The RCS rats, a model for retinal degeneration, displayed normal OMR performance at postnatal days 30 (P30) and 50 (P50), but a significant decline was observed at 3 months. In most rats aged 6 and 12 months, the percentage of correct responses (PoC) was 50%, indicating blindness. The high efficiency and reproducibility of our non-invasive OMR test, along with its suitability for post-surgical and retinal degeneration models, underscores its potential for broad applications in visual performance evaluation.

## 1 Introduction

Rodents, as nocturnal mammals, have vision specialized for detecting motion and contrast in low-light conditions. Despite this, laboratory rodents are capable of complex visual tasks, such as perceiving subjective contours and recognizing visual objects invariantly (1). Vision decline can significantly impact their performance in various behavioral tasks. Numerous rodent models of eye disease have been described and are used for basic research and for testing new therapeutic approaches. Among them are the Royal College of Surgeons (RCS) rats (2), which are a well established model for photoreceptor degeneration and mimic several aspects of retinitis pigmentosa, an inherited retinal degeneration disease, and age-related macular degeneration (3). In this animal model, a defect in the *MERTK* gene leads to a phagocytosis defect in the cells of the retinal pigment epithelium (RPE) and an accumulation of shed outer photoreceptor segments (4). This inability leads to the progressive death of photoreceptor cells (PR) and to visual impairment and blindness within 6 to 12 months (5).

To characterize rodent models of eye disease, it is essential to evaluate the visual function of both diseased and healthy rodents to assess disease progression. By employing a robust test to quantify visual function, the rat models can be utilized to test the efficacy of new therapies.

Rodent vision can be tested through various methods, including training tasks, like visual water maze tasks, reflex-based behavioral tasks, such as optomotor response (OMR) and optokinetic response (OKR), looming response, or electrophysiological recordings. The OMR test, which involves observing an animal’s instinctive head movements to stabilize the retinal image when exposed to a moving grating stimulus (6), is particularly popular due to its non-invasive nature and the lack of required training. The OMR discriminates from the OKR in that the former involves compensatory body movements, i.e. mainly of the head, while in the latter the compensatory movements are made only by the eyes in order to stabilize the image (6).

The OMR test is widely used to assess rodent visual behavior (7). Commercial OMR test systems, such as OptoDrum (Striatech GmbH, Germany), OptoMotry HD (CerebralMechanics Inc., Canada), and qOMR (PhenoSys GmbH, Berlin, Germany), automate the output of visual contrast thresholds, visual acuity, or OMR indices and are commonly used in rodent visual tests (8–10). However, these commercial systems are limited to specific visual stimuli, such as black-white gratings with pre-designed spatial frequencies and drifting velocities.

Optimal spatial frequency and drifting speed of gratings may vary between rodent strains. With the advent of open-source behavioral test resources like DeepLabCut (11, 12), which trains neural networks to track animal positions, and Psychopy (13, 14), which designs visual stimuli for OMR tests, there is an opportunity to develop more customizable setups.

In this study, we have implemented a novel setup for OMR visual tests using DeepLabCut and Psychopy, coupled with our customized algorithm (15). Using pigmented RCS rats and wild-type (WT) controls, we demonstrate the potential of the setup for evaluating the visual performance of rats during the process of vision loss.

## 2 Methods and Materials

### 2.1 Animals

RCS rats (RCS-p+/LavRrrc) and WT rats (RCS-rdy+p+/LavRrrc) were bred locally from four breeding pairs obtained from the Rat Resource & Research Center (RRRC, USA). The animals were housed in an animal facility with a 12-hour light/dark cycle in Blue Line Sealsafe 1500U cages (2–4 rats per cage). They had *ad libitum* access to standard rat chow and water throughout the study.

We used three 5 to 6-month-old WT rats to determine the optimal spatial frequency (SF) and velocity of moving gratings and three 6 to 7-month-old RCS rats for comparison. Subsequently, we divided additional fifteen RCS and fifteen WT rats into five groups with increasing age (P30, P50, 3M, 6M, 12M). Originally designed as a pilot study, each group consisted of three animals for OMR testing at 10 Lux with contrast levels of 1, 0.5, and 0.1. Age and gender of these animals are given in table 1.

**Table 1:** RCS and WT rats used for OMR testing between P30 and 12M.

| group | ID | RCS |  | WT |  |
| --- | --- | --- | --- | --- | --- |
|  |  | Age | Gender | Age | Gender |
| **P30** | 1 | 32d | *M* | 32d | *M* |
|  | 2 | 31d | *M* | 33d | *M* |
|  | 3 | 32d | *F* | 33d | *F* |
| **P50** | 4 | 53d | *M* | 54d | *M* |
|  | 5 | 53d | *M* | 54d | *M* |
|  | 6 | 52d | *F* | 55d | *F* |
| **3M** | 7 | 3m4d | *M* | 3m | *M* |
|  | 8 | 3m2d | *M* | 2m29d | *F* |
|  | 9 | 3m2d | *F* | 2m29d | *F* |
| **6M** | 10 | 6m2d | *M* | 6m5d | *M* |
|  | 11 | 6m2d | *F* | 5m29d | *F* |
|  | 12 | 6m5d | *F* | 6m | *F* |
| **12M** | 13 | 12m10d | *M* | 12m1d | *M* |
|  | 14 | 12m10d | *M* | 11m29d | *F* |
|  | 15 | 12m15d | *F* | 12m | *F* |

Abbreviations: m, month; d, day; M, male; F, female

All animal experiments were approved by the local State Office for Nature, Environment, and Consumer Protection of North-Rhine Westphalia (Landesamt für Natur, Umwelt und Verbraucherschutz Nordrhein-Westfalen, Germany) (81-02.04.2021.A111). The experiments adhered to ARRIVE guidelines (16), the German Animal Welfare Act, and EU Directive 2010/63/EU for animal experiments.

### 2.2 OMR box hardware

The OMR setup for RCS rats was adapted from a mouse OMR box (15, 17) (Fig.1). The rat-specific OMR box was modified in size and featured four 24” monitors (EIZO) mounted onto a custom-built PVC box (530 mm x 530 mm, height 450 mm) with a movable lid and a centrally positioned elevated circular platform. These larger monitors provided ample space for the rats, preventing unintended screen contact. The PVC box was 15 cm higher than the monitors, ensuring sufficient overhead space for comprehensive video capture and preventing interference with the top camera.

**Fig. 1.**
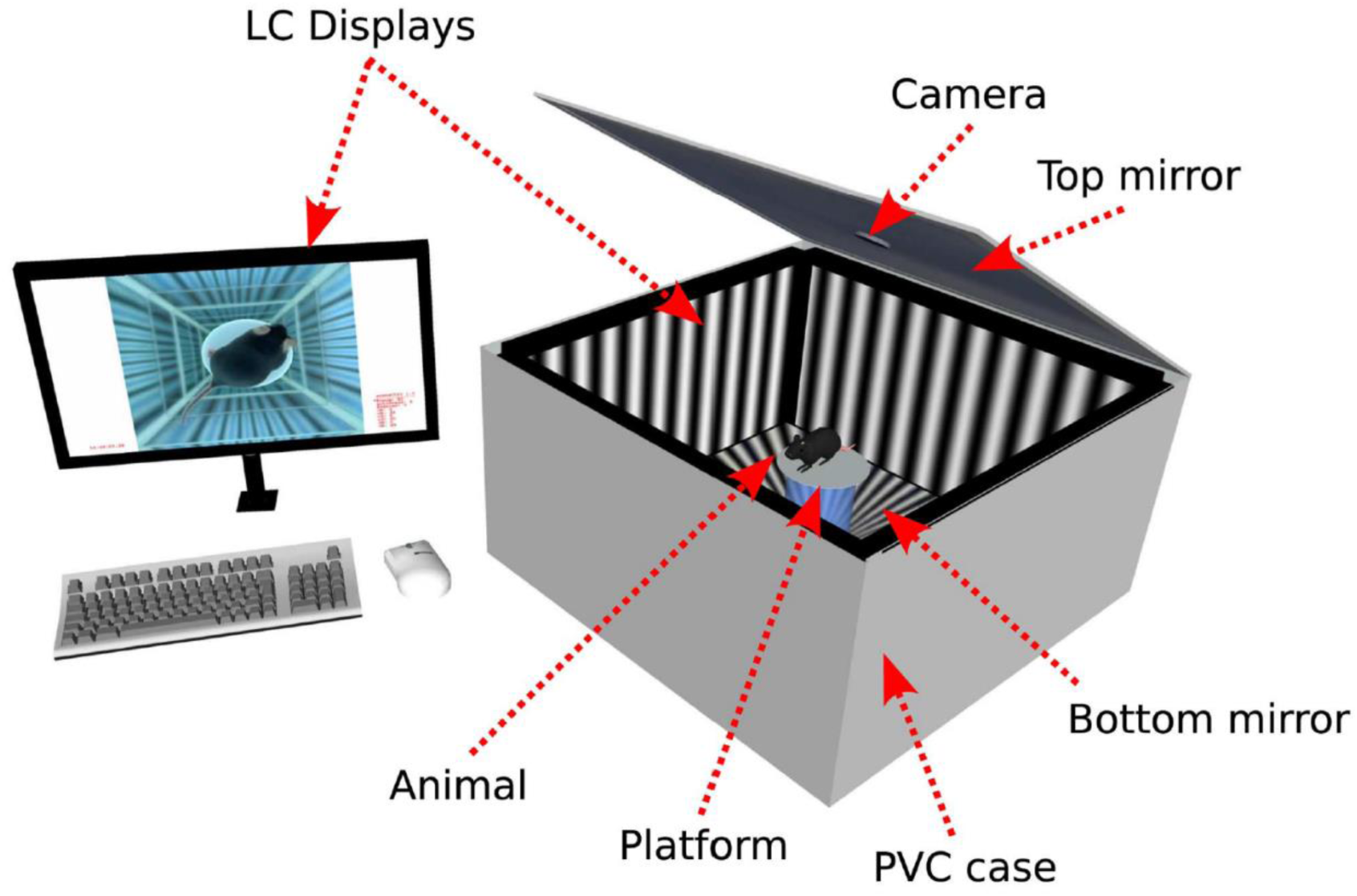
Scheme for a homemade rat OMR box adapted from a previously described mouse OMR box. Four monitors were securely affixed within a PVC box. Reflective surfaces were positioned at the top and bottom to generate an impression of boundless depth, preventing the animal from jumping outside the platform. An overhead camera was installed above the platform to monitor head movement. The experimenter can access the stimulus parameters and head-tracking data via a fifth monitor (image taken from Kretschmer et al. (2013), PLOS One, (CC BY 4.0) (17)).

The 3D-printed PVC plastic circular platform, with a diameter of 14 cm, allowed rats to remain comfortable while restricting random movements. PVC mirrors were strategically placed in three areas to enhance depth perception and deter jumping behavior: the interior area of the box, the bottom floor for ease of cleaning, and the inner side of the lid.

The head-tracking camera (Logitech QuickCam Pro 9000) was positioned at the center of the lid and its autofocus and infrared capabilities were used to capture high-resolution video in low-light conditions. Adhesive foil covered the monitor borders to prevent light from entering the box. The four monitors were connected to a dedicated graphics card (ASUS ROG-STRIX-RTX3070) via an HDMI splitter, with a fifth monitor connected to the PC mainboard (ASUS PRIME Z590-P) for experiment control.

### 2.3 Test optimal SF and velocity of the gratings for RCS and WT rats

Rats, being nocturnal, have photoreceptors adapted to dim lighting conditions (18). For this study, a light intensity of 10 Lux was tested, as higher light levels can cause avoidance behavior and affect head movements during OMR testing (19). The screen brightness and an additional adjustment via Dimmer software (v 2.0.0.b9, Nelson Pires) were set to achieve 10 Lux inside the box, as measured with a luxmeter (PCE-172, PCE Deutschland GmbH) placed at the platform center. Visual stimuli, including contrasts, duration, and direction, were created using PsychoPy software (20) as sequences of clockwise and anticlockwise black-white gratings (Fig. 2A). The spatial frequency refers to the thickness of the bars, while the contrast refers to the difference between the black and white. No gamma correction was applied to the displays. The habituation phase for the animals involved three steps prior to testing: 3 days of housing room adaptation, 30 minutes of acclimatization to the OMR room, and a single 6-minute session of stimuli adaptation. The OMR test was conducted in the morning between 8 a.m. and 12 a.m.

**Fig.2.**
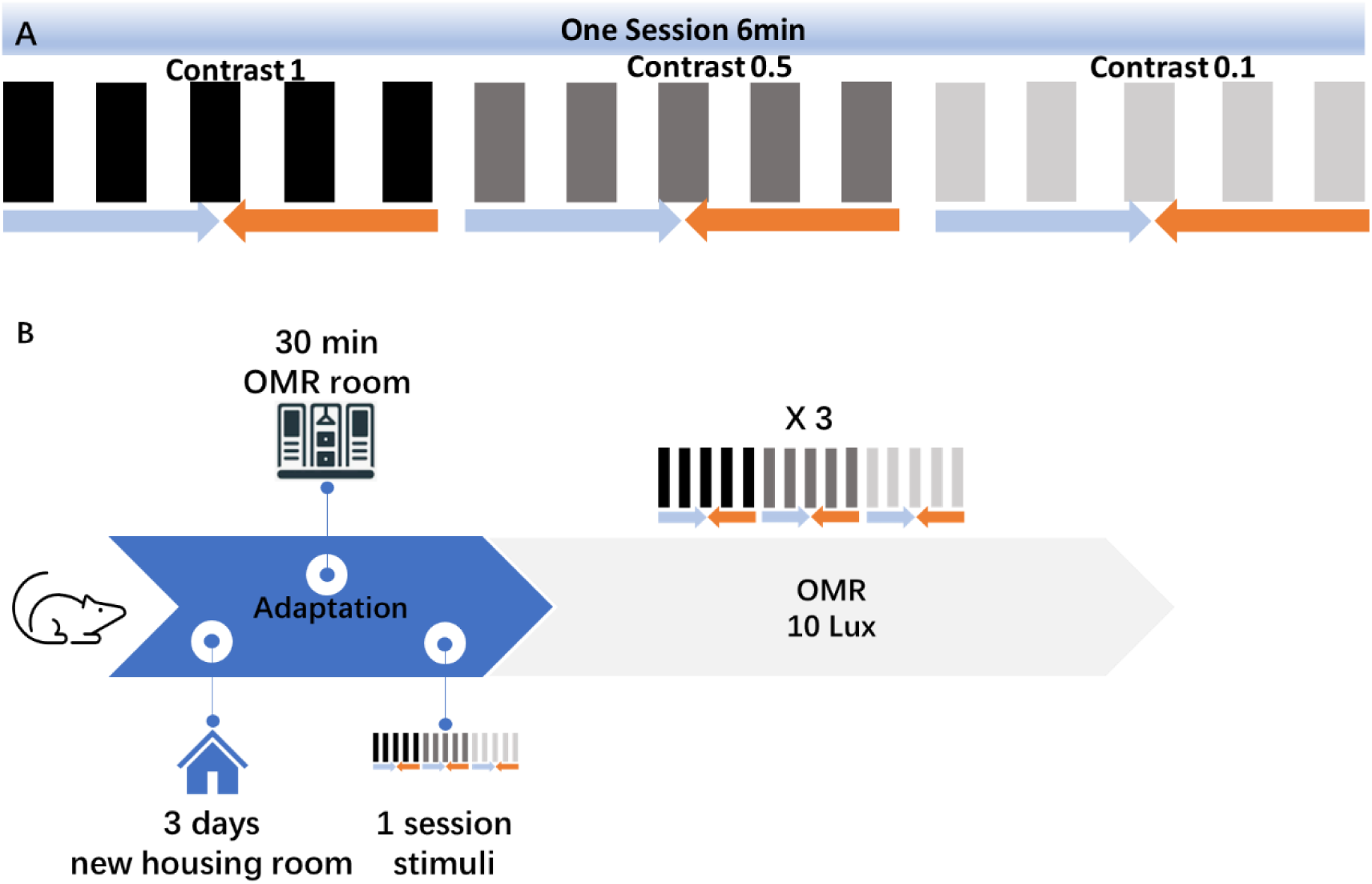
Overview of OMR testing protocol. **(A)** Within one session of 6 min, consecutive gratings varied from contrast 1, to contrast 0.5, to contrast 0.1, which was designed by PsychoPy (20). Gratings moved in one direction for one minute and then changed direction for another minute. **(B)** Rats were transferred from the animal facility to the new housing room 3 days before testing (adaptation phase). On the day of the OMR test, animals stayed in the OMR room for 30 min (acclimatization to the room) followed by one session of grating stimuli adaptation inside the OMR box before three test sessions were started.

Four drifting velocities (0.5 Hz, 1.0 Hz, 1.5 Hz, 2.0 Hz, corresponding to 180°/s, 360°/s, 540°/s, 720°/s, respectively) combined with four spatial frequencies from 0.2 to 0.5 cycles per degree (cpd) (0.2 cpd, 0.3 cpd, 0.4 cpd, 0.5 cpd) were tested at contrast 1 to determine the optimal parameters for RCS and WT rats.

### 2.4 OMR test at 10 Lux

OMR responses were evaluated at 10 Lux using the selected drifting velocity and spatial frequency with RCS rats aged between P30 and 12 months, representing different stages of retinal degeneration. Age-matched WT rats served as control groups, with three rats per group. Each rat underwent three consecutive 6-minute sessions of visual stimuli at 10 Lux, totaling 18 minutes of testing (Fig. 2B). Video recordings were made at 25 frames per second (FPS) from the start of the stimuli.

### 2.5 OMR output analysis

DeepLabCut (11, 12), was used to track the real-time positions of the rats’ ears in the recorded videos, employing a pre-trained neural network (Fig. 3A). The network was trained following a described protocol (11), with outputs including videos in which ears were marked, and the coordinates of both ears (left ear X1 Y1, right ear X2 Y2) were saved in a CSV file (Fig. 3B). Likelihood over 95% was deemed acceptable accuracy, with over 95% of frames meeting this criterion. Frames below this threshold were adjusted manually and used to train a feedback loop in the neural network.

**Fig. 3.**
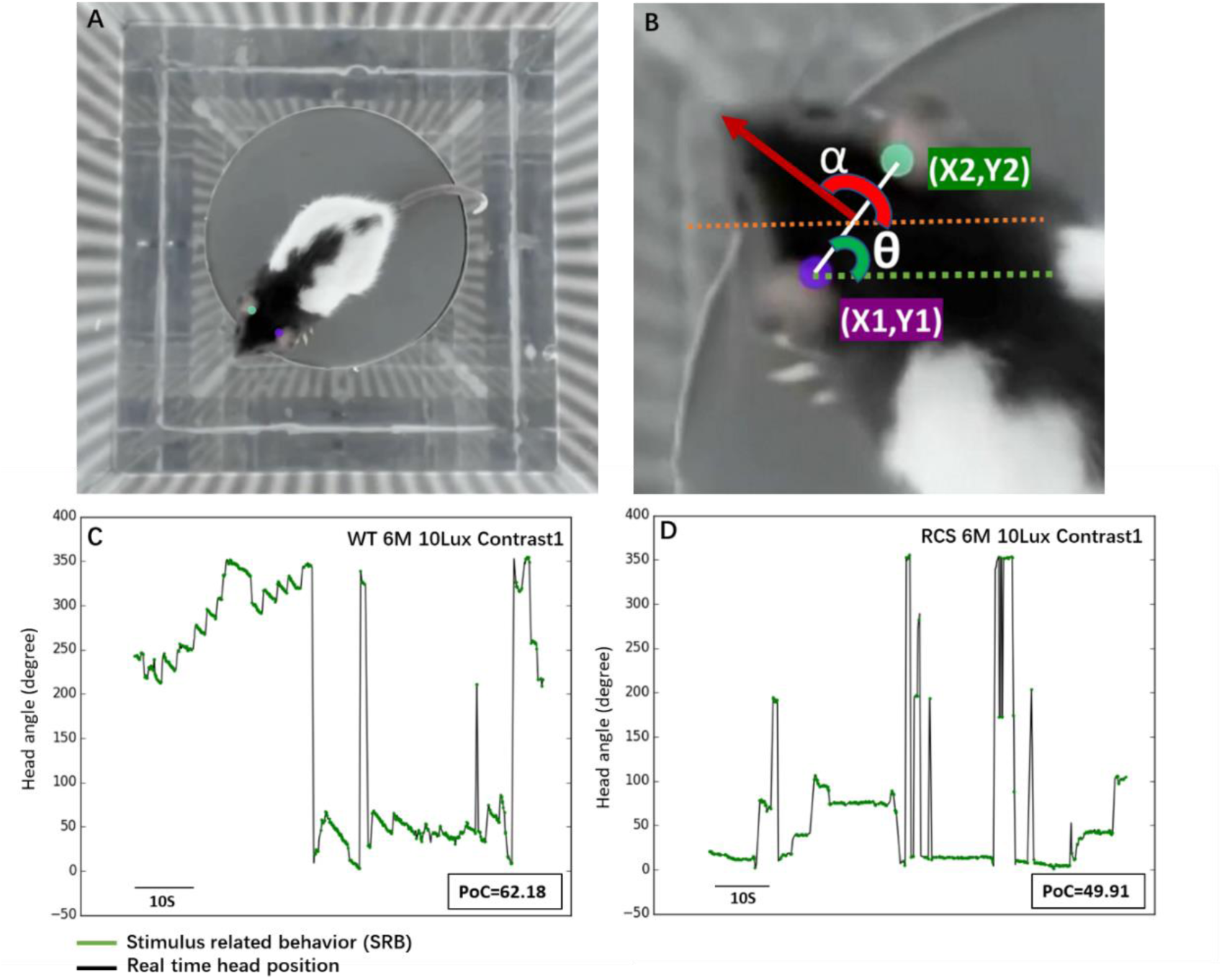
Illustration of PoC analysis. The PoC analysis protocol with an example from a 6-month-old WT and an RCS rat tested at contrast 1 and 10 Lux. **(A)** Illustrative frame extracted from recorded videos marking the recognition of both ears (left ear in purple, right ear in green). **(B)** Display of the real-time head position α, calculated from θ, the angle of the ear line. **(C, D)** The real-time head angle recordings of a 6-month-old WT rat (C) and a 6-month-old RCS rat (D) during the OMR test conducted at contrast 1 and 10 Lux. The black line represents the total frames recorded, while the green line indicates stimulus-related behavior from which the percentage of correct (PoC) responses is calculated.

Using ear coordinates and our customized algorithm, the angle θ between the ears and the x-axis (0 to 90°) was calculated, followed by the real-time head angle α (0 to 360°) (Fig. 3B). With a video frame rate of 25 FPS, the time interval between frames is 1/25 second.

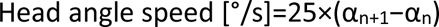

Stimulus-related behavior (SRB) was defined as head movements aligning with the grating direction and falling within a specific velocity range, excluding sudden high-speed movements and minor low-speed adjustments. Previous mouse OMR studies have typically used head velocities between 2°/s and 12°/s (6, 17). However, as rats display more vigorous head movements, velocities between 2°/s to 40°/s were applied in this study.

The percentage of correct (PoC) index was calculated as the proportion of frames displaying SRB out of the total recorded frames. PoCs of 50 are considered indicative of blindness, as the animals’ responses align with chance levels.

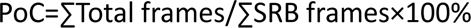

### 2.6 Immunohistochemistry of retinae

Retinae from one rat per group were harvested and processed for immunohistochemistry following described protocols (21). Eyes were enucleated, and retinae in the eye cup were immersion-fixed in 4% paraformaldehyde (PFA) in 0.1 M phosphate buffer (PBS) for 30 minutes at room temperature. The tissue was incubated in 10% sucrose in PBS for 1 hour, followed by 30% sucrose overnight. The retinae were frozen in OCT compound, and vertical sections (perpendicular to the retinal layers, 20 μm) were taken in a cryostat and incubated with primary and secondary antibodies (Table 2).

**Table 2:** Antibodies used for retina staining.

| **Primary Ab** | **Dilution** | **Secondary Ab** | **Dilution** |
| --- | --- | --- | --- |
| rb Recoverin<br>(Chemicon, ab5585-l) | 1/2000 | gt α rb Alexa Fluor 488<br>(Invitrogen, A11034) | 1/500 |
| ms Rhodopsin<br>(Invitrogen, p21940) | 1/500 | d α ms Cy5<br>(Dianova, 715-175-151) | 1/500 |
| rb M-opsin<br>(Chemicon, ab5405) | 1/2000 | d α rb Cy2<br>(Dianova, 711-225-152) | 1/400 |
| gt S-opsin<br>(Santa Cruz, sc-14365 ) | 1/200 | d α gt Cy3<br>(Dianova, 705-165-147) | 1/1000 |

Abbreviations: Ab, antibody; gt, goat; ms, mouse; rb, rabbit; d, donkey

Whole-mount retinae were also incubated with primary antibodies for 3 days, followed by secondary antibodies overnight. The stained retinae were examined using a confocal laser scanning microscope (TCS SP5 II; Leica Microsystems) with 63x/1.4 oil immersion lenses.

Biotinylated peanut agglutinin (PEA) (1/1600, Sigma, I6135), and streptavidin Alexa Fluor 647 (1/100, Molecular Probes) were used for double staining retina sections and whole-mount retinae, marking inner and outer segments of cones and cone terminal systems in the outer plexiform layer (22).

### 2.7 Statistical analysis

Statistical analysis was conducted using GraphPad Prism 9.5.0. Differences in OMR performance between the RCS and WT rats were evaluated using a Two-Way ANOVA, with Gaussian distribution validated beforehand by normality QQ-plots. Multiple Tuke’s or Sidak’s post-hoc comparisons were applied to determine significance for the optimal spatial frequency and velocity, and differences between WT and RCS rats, respectively. Levels of significance are indicated by asterisks: **** for p ≤ 0.0001, *** for p ≤ 0.001, ** for p ≤ 0.01, and * for p ≤ 0.05. All values are presented as mean ± standard error of the mean (SEM).

## 3. Results

### 3.1 PoC analysis protocol with examples

Figure 3 illustrates the PoC analysis protocol with examples from a 6-month-old WT rat and an RCS rat tested at a contrast of 1 and 10 Lux. The ratś ears were recognized by DeepLabCut (Fig. 3A), and their positions were used to determine the real-time head position α, calculated from θ, the angle of the line between the two ears (Fig. 3B). As illustrated in Figs. 3C and 3D, the real-time head angles are plotted over the entire duration of the test session with a 6-month-old WT rat and a 6-month-old RCS rat conducted at contrast 1 and 40 Lux. Stimulus-related behavior (SRB), defined as head movement aligned with the grating direction and head angle speed within the range of 2-40°/s, was recognized by the software, resulting in PoC values of 62.18 and 49.91 for the WT and RCS rat, respectively.

### 3.2 Optimal spatial frequency and velocity for both RCS and WT rats

WT rats aged between 5 and 6 months were used to determine the optimal spatial frequency (SF) and grating velocity in the OMR test. RCS rats aged between 6 and 7 months were used in comparison. At that age, degeneration of the retina is in an advanced stage in RCS rats, and the animals are considered to be blind (2). Figure 4 shows that the RCS rats exhibited no differences in OMR performance across four spatial frequencies (0.2, 0.3, 0.4, 0.5 cpd) and four grating velocities (0.5, 1.0, 1.5, 2.0 Hz). In WT rats, the OMR performance at 0.3 cpd and a velocity of 2 Hz (PoC=66.87±22.05) was significantly higher than at 0.5 cpd and a velocity of 1 Hz (PoC=56.69±3.43, P=0.0156) and 0.5 cpd and a velocity of 1.5 Hz (PoC=57.46±3.39, P=0.034). No differences were observed in OMR performance among other parameters in the WT rats. Since this combination yielded the highest performance of WT rats, 0.2 cpd and 3 Hz were set as the SF and velocity for both the RCS and WT rats in the subsequent OMR tests.

**Fig. 4.**
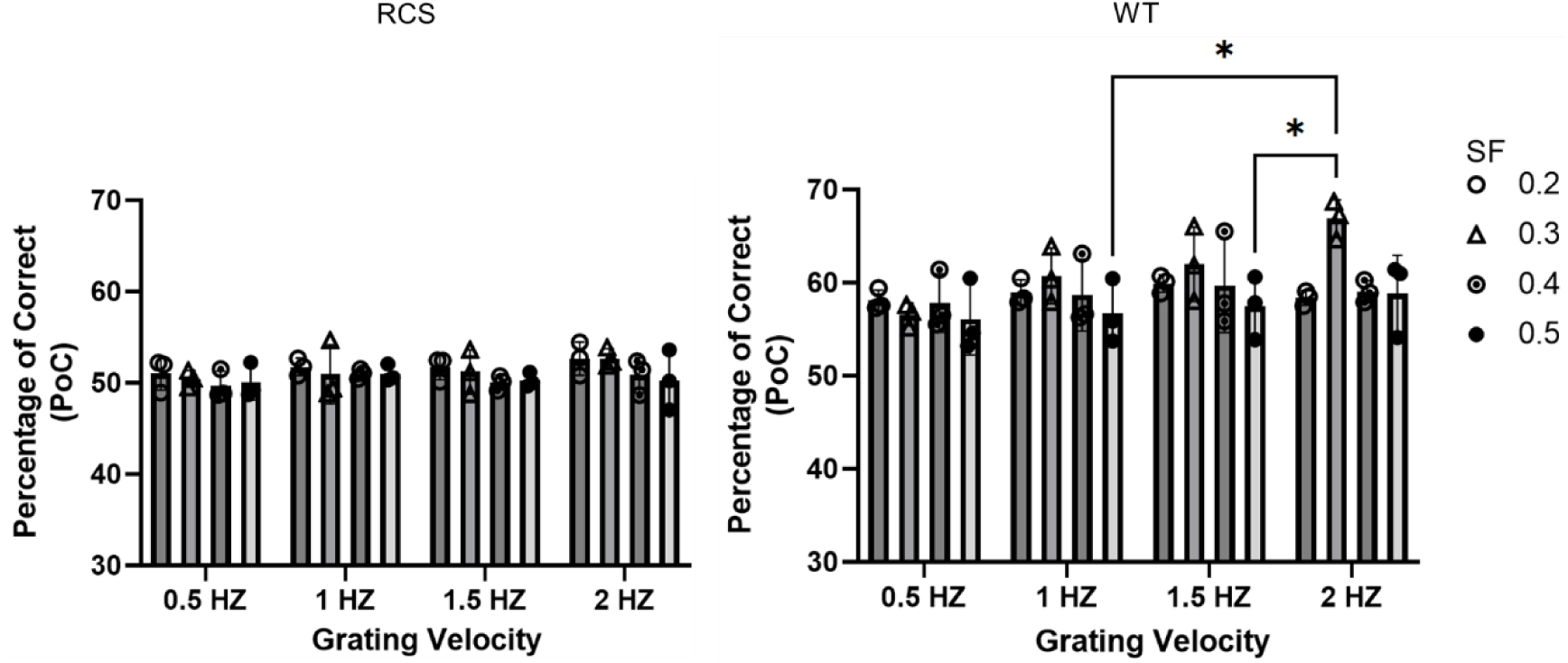
Optimal spatial frequency and velocity for RCS and WT rats. RCS rats (6 to 7 months of age) showed no differences across four spatial frequencies (SF) and four grating velocities. In age-matched WT rats, OMR performance at 0.3 cpd and a velocity of 2 Hz was significantly higher than at other visual stimuli. * p ≤ 0.05

### 3.3 Significant decrease in OMR performance in RCS rats from 3 months onward

To follow the visual performance of the RCS rats during the process of retinal degeneration, OMR measurements were performed with RCS and WT rats at different ages using three contrast levels (high: 1.0, middle: 0.5, low: 0.1) under a light intensity of 10 Lux. As shown in Fig. 5, RCS rats generally maintained normal vision at P30 and P50 compared to age-matched WT rats. At 3 months, RCS rats showed a significant decline in PoC compared to WT rats (contrast 1: PoC_RCS_=56.7±3.2 vs. PoC_WT_=63.6±2.5, p=0.0305; contrast 0.5: PoC_RCS_=57.1±2.8 vs. PoC_WT_=63.4±0.6, p=0.0163; contrast 0.1: PoC_RCS_=51.4±2.1 vs. PoC_WT_=56.3±3.3, p=0.0265). This decline persisted in 6-month and 12-month-old RCS rats, with more substantial differences compared to WT rats as degeneration progressed. As soon as the PoC drops to 50, it can be assumed that the animals are blind, as their response is at chance level. Most of the 6-month and 12-month-old RCS rats were considered blind at all three tested contrasts at 10 Lux light intensity. However, one animal performed above chance level even at the advanced age of 12 months.

**Fig. 5.**
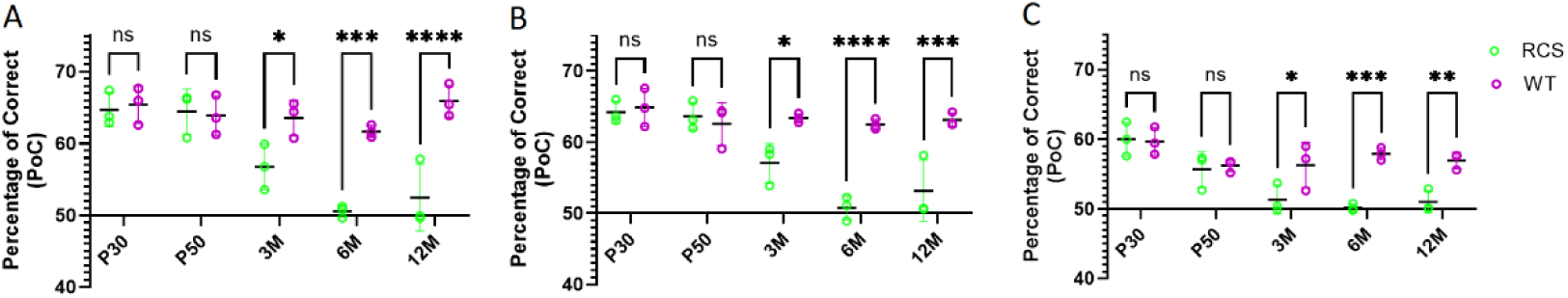
OMR performance in RCS and WT rats upon aging. OMR was tested at 10 lux and three different contrasts (A, 1 = high; B, 0.5 = middle; C, 0.1 = low) with RCS and WT rats from the age of 30 days (P30) until 12 months of age (12M). The RCS rats maintained normal vision at P30 and P50 compared to WT rats. At 3 months, RCS rats showed a significant decline in PoC compared to WT rats, which persisted at 6M and 12M.

### 3.4 Degeneration of photoreceptors, especially rods, in RCS rats

Next, we stained the retinae of RCS rats at different ages for rod and cone photoreceptor markers. Recoverin is a neuronal calcium-binding protein primarily detected in the PRs of the eye. Antibodies against recoverin selectively label both types of PRs, namely the somata in the outer nuclear layer (ONL), the outer segments (OS), and inner segments (IS) of rods and cones, and also identify two types of cone bipolar cells (BCs) in the rat retina (23, 24). Additionally, an antibody targeting rhodopsin was used to stain rod OS, as the photopigment rhodopsin is exclusively present in rods.

In Fig. 6, the staining against recoverin (green) and rhodopsin (blue) is shown in a representative adult WT retina as well as in different postnatal stages of four RCS retinae (1M, 3M, 6M and 12M). In the WT retina, the PR somata in the ONL, as well as their IS and OS, and their terminal systems in the outer plexiform layer (OPL) were recoverin-positive. The two types of BCs in the inner nuclear layer (INL) and their terminals ramifying in two strata in the inner plexiform layer (IPL) were also recoverin-positive. Rod OS were strongly labeled for rhodopsin. In the WT retina, the ONL contained 10-11 rows of PR somata. Owing to the fact that cones make up less than 5% of photoreceptors in rat retina (25, 26), a nearly continuous band of rod OS was marked by rhodopsin.

**Fig. 6.**
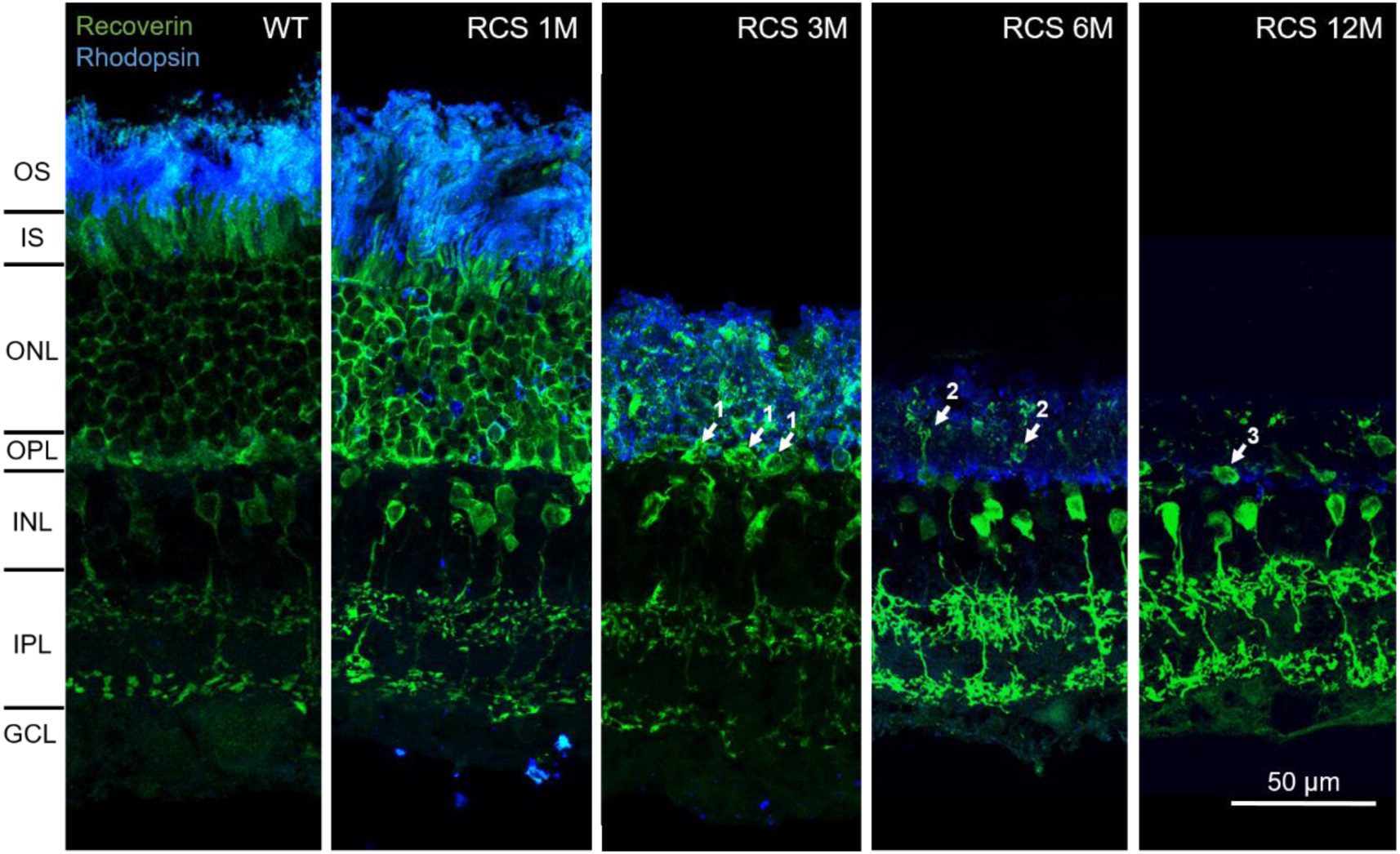
Retinal sections of WT and RCS rats at different postnatal stages stained against recoverin and rhodopsin. In the WT retina, recoverin positivity (green) was observed in the intact ONL, IS, OS, and terminal systems of PRs and in two bipolar cell types. In the RCS retina, PRs were gradually lost. Arrowhead 1: a discontinuous row of PRs; arrowheads 2 and 3: Single surviving recoverin-positive PRs. OS: outer segments, IS: inner segments, ONL: outer nuclear layer, OPL: outer plexiform layer, INL: inner nuclear layer, IPL: inner plexiform layer, GCL: ganglion cell layer.

In the RCS retina at 1 month, the ONL was slightly reduced in thickness. The thick rhodopsin-positive stratum comprises not only OS but also a layer of non-phagocytosed OS (debris). By 3 months, most photoreceptors had disappeared. Only a discontinuous single row of PR somata (Fig. 6, arrowheads 1) remained. A layer of debris consisting of non-phagocytosed outer segment material immunoreactive for recoverin and rhodopsin was observed.

At 6 months, the debris zone was thinner. Debris was rhodopsin-positive. Recoverin immunoreactivity was found in somata and thin structures (arrowheads 2), most likely representing processes grown by photoreceptors. At 12 months, debris was further reduced. Some recoverin-positive somata were found that might represent surviving photoreceptors (arrowhead 3).

In summary, a reduction of the ONL was evident in the RCS retinae from 1 to 3 months of age. In the later stages (6 and 12 months), isolated recoverin-positive somata, most likely PRs, were detectable in what had remained of the outer RCS retina; however, uncertainty remains as to whether those surviving PRs were cones.

### 3.5 Remaining photoreceptors in older RCS rats are cones

Since the recoverin antibody does not differentiate between rods and cones, given the proteińs presence in both cell types, specific antibodies targeting the two opsins exclusively to cones were utilized to specifically stain these PRs. In Fig. 7, the long-wavelength-sensitive (red/green) opsin and the short-wavelength-sensitive (blue) opsin were used to stain the OS of cones in the WT and RCS retinae. Notably, in pigmented rat strains, only around 9% to 10% of the cone population is formed by blue cones, while the majority are red/green cones (27). Fluorescently marked peanut lectin (PEA) (appearing blue in Fig. 7), was employed to label all cone IS, OS, and cone terminal systems in the OPL (22).

**Fig. 7.**
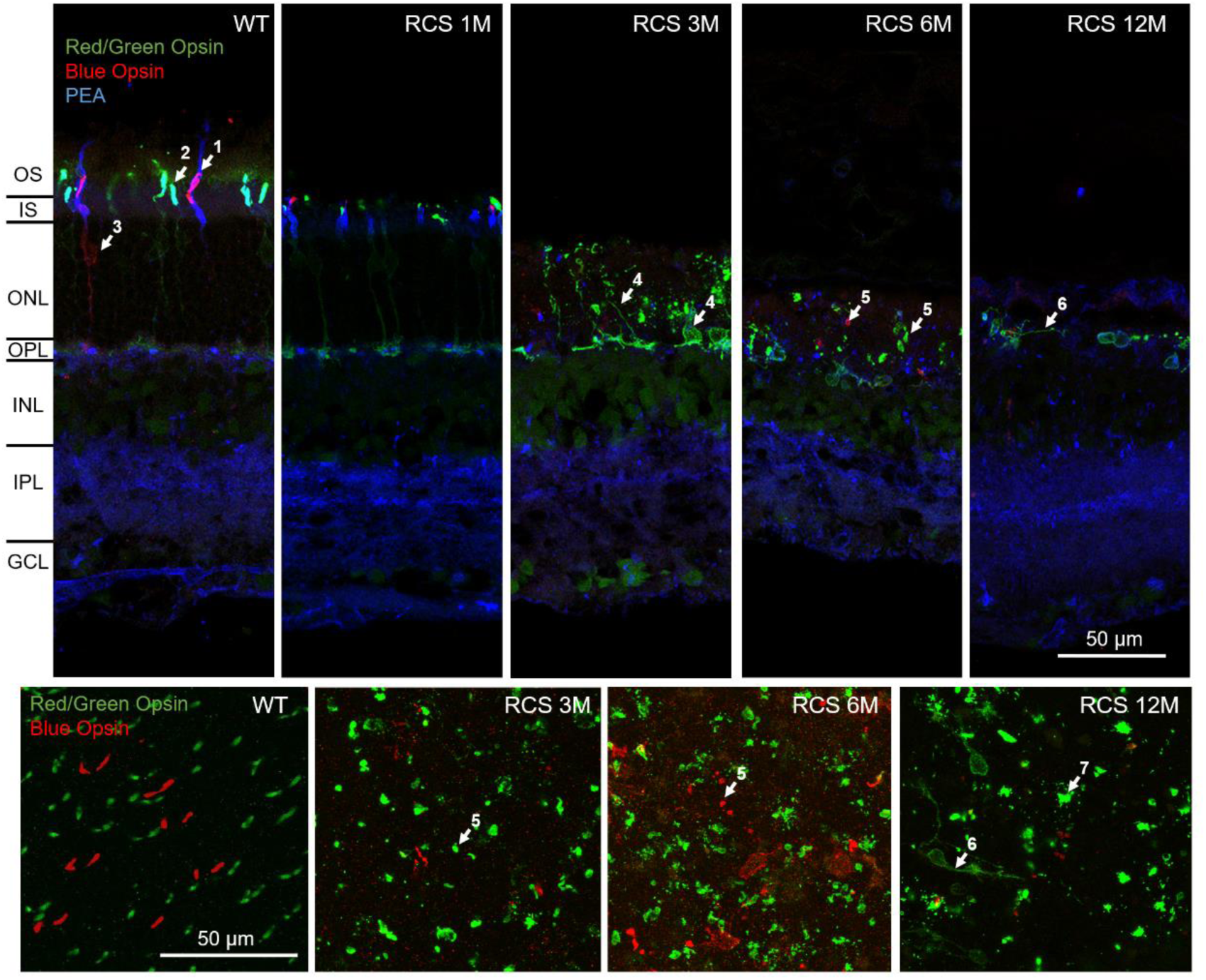
The remaining PRs in the late stage of retinal degeneration are cones. Upper row, staining of cross-sectional retina sections with antibodies against red/green opsin (green), blue opsin (red) and peanut lectin (PEA, blue, labeling all cones). Arrowhead 1: purple color of cone OS expressing blue opsin counterstained with PEA; arrowhead 2: cyan color of red/green opsin-positive OS counterstained with PEA; arrowhead 3: only occasionally, somata and axons were weakly labeled by the opsin antibodies in WT. Compared to the WT retina, cone OS were considerably shortened in the RCS retina at 1 month (1M). From 3 (3M) to 12 months (12M), cone degeneration was evident, with OS becoming shorter and disorganized and opsins being expressed throughout the entire cell. Arrowhead 4: cone soma and processes running through the debris layer are opsin-positive. Arrowhead 5: short or fusiform cone OS. Arrowhead 6: processes running horizontally. Arrowhead 7: large structures are most likely detached cone OS. OS: outer segments, IS: inner segments, ONL: outer nuclear layer, OPL: outer plexiform layer, INL: inner nuclear layer, IPL: inner plexiform layer, GCL: ganglion cell layer.

In Fig. 7, within the WT retina, three distinct purple-appearing OS (marked by arrowhead 1 in Fig. 7) were identified, indicating the presence of blue opsins (appearing red) counterstained with PEA (appearing blue). Red/green opsins expressing OS (arrowhead 2 in Fig. 7) exhibited a cyan hue. Note that somata and the axons of cones were only occasionally weakly labeled by the opsin antibodies (arrowhead 3 in Fig. 7).

In the retinae of RCS rats at one month of age, cone opsins in OS exhibited a reduction to less than 50% of their normal length. In later stages, cones became more and more disorganized. Opsin immunoreactivity could be found throughout the entire PR, including soma and axon (arrowheads 4 in Fig. 7). Cone OS were strongly reduced, sometimes appearing as dots or spindle-shaped (arrowheads 5 in Fig. 7). The rudimentary OS might explain the residual vision observed in RCS rats at the age of 3 months. At 6 and 12 months, opsin-immunoreactive somata and processes (arrowheads 6), as well as large malformed recoverin-positive structures, most likely fragments of detached and non-phagocytosed cone OS (arrowhead 7), were observed. While the opsin-immunoreactivity identifies the cells as cones, the lack of a clear OS connected to the somata and the numerous horizontal processes show that the cells have gone through a considerable process of degeneration and restructuring.

To summarize, the RCS retinae at 3 months exhibited preserved cone structures with shortened OS. At the age of 6 and 12 months, the regular cone patterns vanished, although several cone-like cells were still identifiable, albeit with abnormal morphology.

## 4. Discussion

In this study, we established an optimized OMR box for rats modeled on existing commercial OMR setups. We implemented a novel method combining DeepLabCut, PsychoPy, and customized algorithms to automate the evaluation of the OMR performance in tested rats (15). By applying the new setup, we determined the optimal SF and grating velocity for the pigmented RCS strain in comparison to WT rats. We were able to show that RCS rats maintained normal OMR performance up to the age of 50 days. After three months, however, a significant decline in OMR performance was observed in different illumination conditions, which became stronger after 6 and 12 months. At this point, most of the RCS rats appeared to be effectively blind, exhibiting a PoC of around 50 %.

### 4.1 Efficiency of OMR test

These results demonstrate that the OMR setup presented here is suited for the detection and tracking of progressive retinal deficits and loss of vision in RCS rats. The advantage of a tailor-made system over commercial systems is, in addition to reduced costs, the flexibility to be able to adjust all parameters. For example, it is necessary to redetermine the optimal baseline spatial frequency and contrast sensitivity with each rat line, as not all lines have the same baseline levels. Brown-Norway rats, for instance, exhibit a higher baseline spatial frequency than Long-Evans rats (28). The implementation of automatic scoring, which reduces the experimenter bias, is also advantageous. The PoC read-out was robust, demonstrating impaired OMR performance in older RCS rats, and significance was reached with only three animals per group. Since the animals were unrestrained, the relative lack of stress may have led to more consistent behavior. Drifting gratings did attract the attention of tested animals immediately following stimulus onset, and behavior could be elicited simultaneously. The use of customized algorithms enhanced the efficiency of analyzing the results from the recorded videos of animal behavior. A common problem that occurs with OMR measurements is the animal jumping off the platform, especially if measured repeatedly. In the system presented here, the built-in mirrors help to create an illusion of depth, which is intended to decrease the occurrence of jumping. The OMR test is usually performed with drifting gratings. However, more naturalistic visual stimuli, including broadband orientations and spatial frequencies, have been shown to be much more efficient in activating visual neurons in the cortex and superior colliculus (29). The use of a custom-built OMR chamber allows the presentation of different visual stimuli, which would be a promising line of future research.

The significant advantage of the OMR test over alternative visual behavioral tests is its high reproducibility with short measurement duration. Alternative tests, such as visual water tasks (30) or operant conditioning (31) require the animals either to be trained, which is more time-consuming and more stressful for the animal. Visual tests that can be performed without prior training include the visual placing and visual cliff tasks and the light/dark box (31, 32).

### 4.2 OMR lasts relatively longer than photoreceptor viability

A progressive decline of visual abilities was described for the pigmented RCS strain used here (30). Using our OMR setup, the PoC values for pigmented RCS rats remained comparable to WT at P30 and P50 but started to decline at 3 months. At 6 and 12 months, correct head-following-gratings behavior was undetectable in most of the rats, with PoC near 50%, i.e., chance level. A similar time course for OMR performance was also described in the literature, although other parameters were selected as read-out (33), showing validity of the new set-up. These findings are comparable to recent reports in *rd10* mice, a mouse model of retinitis pigmentosa, which show complete blindness after at least 6 months (15).

How does the time course of OMR performance match photoreceptor degeneration as studied by histology and electrophysiology? By P50, the outer nuclear layer is consistently degenerated, leaving 4-5 layers of photoreceptors, compared to 11-12 layers in WT (34). At this age, the OMR performance of the RCS rats was indistinguishable from that of WT rats. It seems that the remaining photoreceptors were sufficient to provide an OMR response. By 3 months, only a discontinuous layer of photoreceptors remained in the RCS retinae, and animals showed the first signs of deficits in OMR performance. At 6 and 12 months, only scattered cone-like cells expressing red/green opsin with malformed or lost OS were evident, and the visual function of the animals was lost.

Previous ERG evaluations in pigmented RCS rats have shown fairly normal b-wave (activity mostly attributed to bipolar cells) and scotopic threshold responses up to day 27, with a-wave (photoreceptor response) rarely recordable by day 50 even at maximum intensity, and b-wave not detectable by 100 days (2, 35). The ERG is a well-established method; however, as the recording is performed on the entire eye, small photoreceptor responses that may be sufficient to trigger OMR responses may remain undetected. However, certain discrepancies between histological and functional studies have been reported in the literature. A decrease in the visual performance of pigmented RCS rats at 6 months of age was reported using a water escape paradigm (30), while a reduction in photoreceptor cells was found to follow a different time course. Inconsistencies between histological and functional read-outs have been observed in other rat retinitis pigmentosa models, such as S334ter-3 mutant rhodopsin rats and P23H rats (36), and one study even suggests that a diminished OKT was still detectable in RCS rats even at 10 months (37).

The discrepancies found in the literature suggest that functional performance may be affected by several variables. While it was beyond the scope of our study to solve these discrepancies, our data suggest that surviving cones in 3-month-old RCS rats may account for preserved OMR performance. At 6 and 12 months, cone opsins were likely nonfunctional with malformed and detached OS, supporting the premise that the RCS rats were blind, although single animals were still able to react to the OMR signal.

### 4.3 Conclusion

In summary, in this study, an OMR setup for rats was successfully optimized. A combination of DeepLabCut, PsychoPy, and custom algorithms was used to automatically determine OMR performance in rats. The system demonstrated cost efficiency, flexibility, and high reproducibility with minimal stress for the animals. Validating this setup, we monitored the deterioration of OMR performance in the RCS rats with increasing age compared to the WT rats. Our results indicate a residual function of cones, even after structural reorganization of the retina at 3 months of age, and illustrate the complexity of correlating visual function with histological data. The role of cones in later stages of retinal degeneration requires future investigation.

## 4.5 Acknowledgements

We thank Christoph Aretzweiler von Schwartzenberg for his excellent technical assistance. Special appreciation goes to the developers of DeepLabCut and PsychoPy for providing the tools necessary for our study.

